# Experimental Evidence for Constraints in Amplitude-Timescale Co-variation of a Biomolecular Pulse Generating Circuit Design

**DOI:** 10.1101/756049

**Authors:** Abhilash Patel, Shaunak Sen

## Abstract

Understanding constraints on the functional properties of biomolecular circuit dynamics, such as the variation of amplitude and timescale of pulse, is an important part of biomolecular circuit design. While the amplitude-timescale co-variations of the pulse in an incoherent feedforward loop have been investigated computationally using mathematical models, experimental support for such constraints is relatively unclear. Here, we address this using experimental measurements of an existing pulse generating incoherent feedforward loop circuit realization in the context of a standard mathematical model. We characterize the trends of co-variation in the pulse amplitude and rise time computationally by randomly exploring the parameter space. We experimentally measured the co-variation by varying inducers and found that larger amplitude pulses have slower rise time. We discuss the gap between the experimental measurements and predictions of the standard model, highlighting model additions and other biological factors that might bridge the gap.

## Introduction

An examination of the limits to which important functional properties can be varied provides a design guide for achievable system performance. Examples include the gain-bandwidth constraint in electronic amplifiers [1], the Cramer-Rao bound in statistics [2], and the space-time constraint in software [3]. Investigations in biology from a systems perspective, particularly in the dynamics of biomolecular circuits, have provided instances of such constraints, for example, in the robustness and efficiency of glycotic oscillations [4], responsiveness to noise susceptibility in yeast galactose network [5], the effectiveness and optimality of generalized homeostasis system [6], and sensitivity and adaptation ability in feedforward loop [7]. These constraints provide a guide to the limits of achievable performance in biomolecular circuit dynamics.

Pulses in protein activity (Fig. 1A), in particular, are important dynamics in biomolecular circuits [8]. Both the amplitude of the pulse as well as its duration may be functionally important, for example, in the timescale-based regulatory activity of *crz1* in yeast proteomes [9] and in the pulse amplitude dependent cellular differentiation in *Bacillus subtilis* [10]. One way to generate pulses is when a step input is applied to an incoherent feedforward loop (Fig. 1B). These are a class of biomolecular circuits that are recurring motifs [11] and have been investigated, both in natural circuits and in synthetic circuits, in the context of adaptation [12], fold-change detection [13], and scale invariance [14]. In the context of adaptation, it has been found, computationally, that the sensitivity to input is constrained to be inversely proportional to the adaptation ability [7]. Variations of these relations have been found in other feedforward loops as well [15]. Finally, we have previously addressed, also computationally, the quantitative co-variation between the amplitude and timescale properties for a standard model and noted the different trends possible [16]. An understanding of such constraints can guide the design space of pulse generating biomolecular circuits.

**Figure 1:**
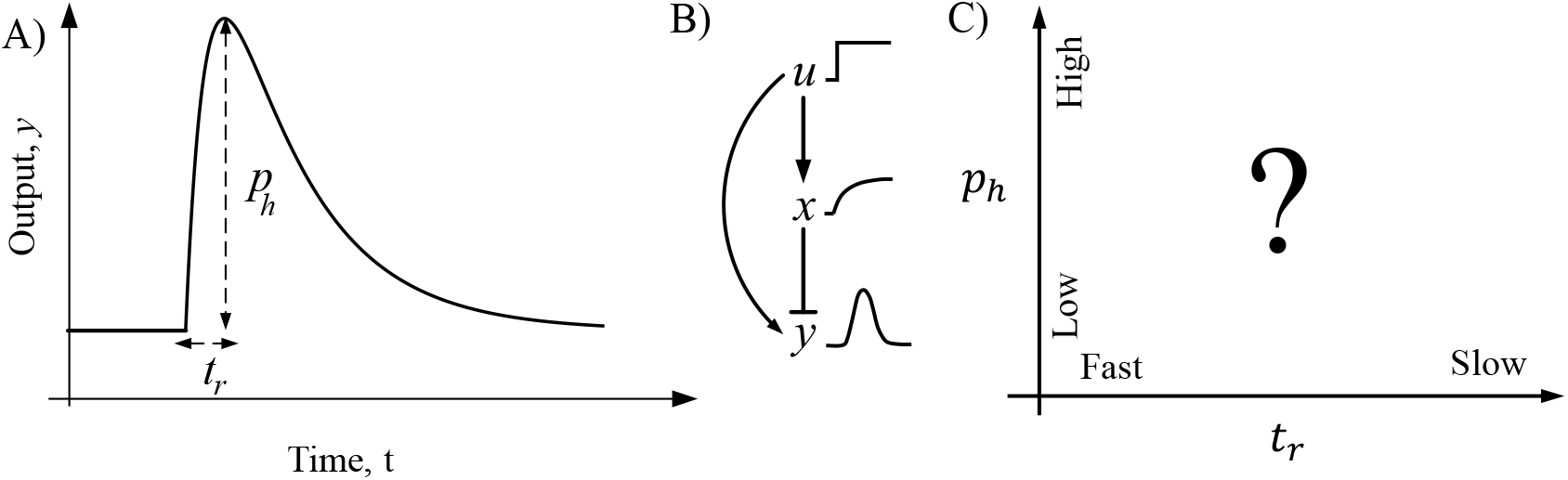
A) Illustration of a pulse. The *p_h_*, and *t_r_* denote the maximum amplitude and the time to reach this amplitude respectively. B) Schematic diagram of an incoherent feedforward loop and typical trajectories of protein concentrations for a step input C) Amplitude-timescale where different pulse trajectories may exist.

There are at least three striking aspects in the dynamics of pulses generated by biomolecular circuits. First, is the wide prevalence of such dynamics, perhaps reflecting the dynamic environments the cells experience, with more instances of pulsing being found due to advances in measurement techniques. Second, that both amplitude and timescale of the pulse may have biological regulatory activity and hence be of functional importance. Third, these amplitude and timescale properties may be interlinked in that they may co-vary in response to changes in circuit parameters, rather than be independently tunable properties. Given these, it is important to experimentally investigate the possible co-variation in amplitude and timescale.

Here we ask whether there is any experimental evidence of constraints in the co-variation of the amplitude and timescale properties of a pulse generating circuit (Fig. 1C). To address this, we used experimental measurements of an existing incoherent feed-forward loop circuit realization and analysed these in the context of a widely accepted mathematical model. We randomly sampled the parameter space of the mathematical model and categorized the co-variations of pulse amplitude and rise time as individual parameters are varied, finding different trends such as mutual increase and mutual decrease. We experimentally characterized these co-variations using inducers and found that as the pulse amplitude increased, the rise time always increased providing evidence for a tradeoff between pulse height and pulse speed. We discuss this disconnect between the model and experiment, investigating model additions and other possible biological factors that might underlie the observed behaviour. These results help to understand the constraints in design of biomolecular pulsing circuits and may also be relevant to naturally occurring biomolecular circuits that pulse.

## Results and Discussion

### Parameter space exploration for amplitude-rise time co-variation

We have previously investigated co-variations of amplitude of pulse with respect to the timescale properties including rise time [16]. We did this using a standard model [17] at a nominal parameter space by individually varying one parameter at a time (see Methods). We found that amplitude and rise time can mutually increase as degradation rate of the output protein increases, and that amplitude can decrease with an increase in rise time as degradation rate of intermediate protein (*x*) increases. Further, we found that only the pulse amplitude, not the timescale properties, change when the production factors change. These are summarized in Fig. 2A.

**Figure 2:**
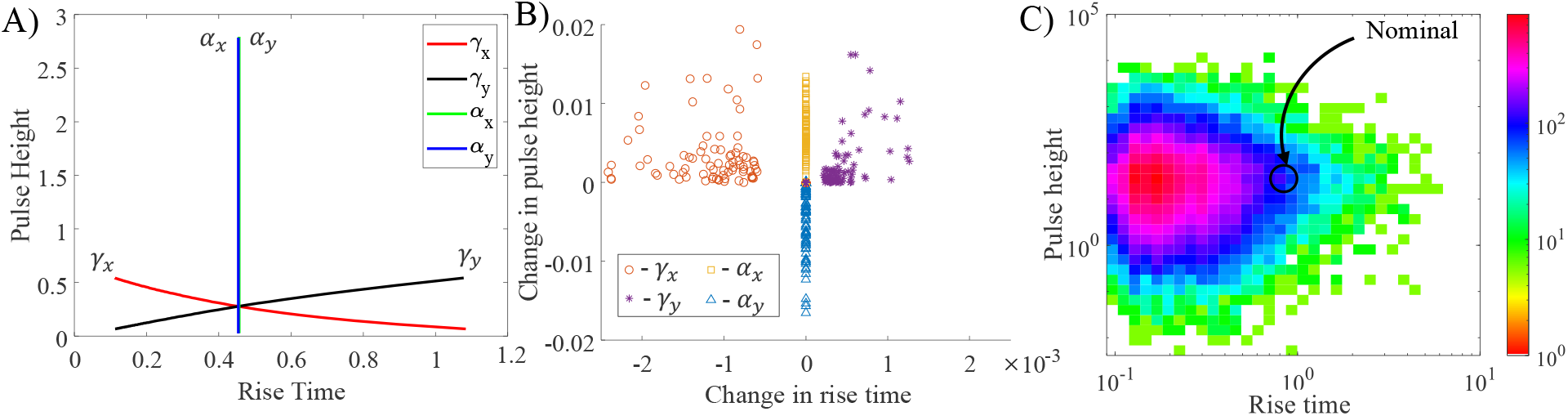
Trends for amplitude-timescale co-variations as pulse height and rise time with parameter space exploration. A) Solid lines are the amplitude and rise time co-variations as the parameters are individually varied for the nominal parameter parameters. B) Symbols represent the change in pulse height and change in rise time as different randomly sampled points in parameter space as individually parameters are perturbed. C) Colorbar represent the density of parameters at a particular point in the amplitude-rise time space as the multiple parameters are simultaneously perturbed. The encircled point shows the nominal parameter set.

To investigate whether these trends persist for other points in the parameter space, we randomly varied the parameters around the nominal parameter set and, for each of the points, computed the change in pulse height and the change in rise time as each parameter is individually perturbed (Fig. 2B). For degradation rate parameter of output protein *y*, the points lie in the first quadrant implying that for an increase in pulse height the rise time also increases. For the degradation rate parameter of intermediate protein *x*, the points lie in the second quadrant implying that an increase in pulse height results in a decrease in rise time. For the production rate of the proteins, the points lie on the y-axis showing that the amplitude can increase or decrease without altering the rise time. Therefore, the trends noted earlier persist across these parameter sets as well.

To further study the effect of perturbations to simultaneous multiple parameters, we randomly perturbed the parameters. The parametric density plot on the amplitude and rise time space shows a larger density of relatively lower amplitude and faster rise time (Fig. 2 C).

In summary, we note that for this model, the amplitude of the pulse height can independently be varied without altering rise time on perturbing production factor of proteins. The amplitude and rise time vary inversely when degradation rate parameter of intermediate protein is changed and they vary proportionally when degradation rate parameter of output protein is changed.

### Experimental Evidence of Co-variation

We obtained a previously constructed incoherent feedforward loop circuit (Fig. 3A inset, Please see Materials) that pulses in response to a step change in *arabinose* in presence of *anhydrotetracycline (aTc)*. In this circuit, the transcriptional activator *AraC* is constitutively produced. The transcriptional repressor *TetR* is expressed from a *AraC*-regulated promoter *P_BAD_*, and expression is activated when inducer *arabinose* is added. A degradation tagged green fluorescent protein (*deGFPssrA*) is expressed from a combinatorially regulated promoter; repressed by *TetR* and activated by *AraC* (in presence of *arabinose*). This is the output of the circuit. The input is a step change in *arabinose*. It is reported previously that *TetR* dominates activation in this circuit [18]. *aTc* binds to and inactivates *TetR*, thereby weakening repression. We replicated the pulse with *arabinose* step in the presence of *aTc* (Fig. 3A, Methods). We find a pulse height of 2600 fluorescence unit and rise time of 75 minutes for 0.2 % *arabinose* and 1 *ng/ml aTc*.

**Figure 3:**
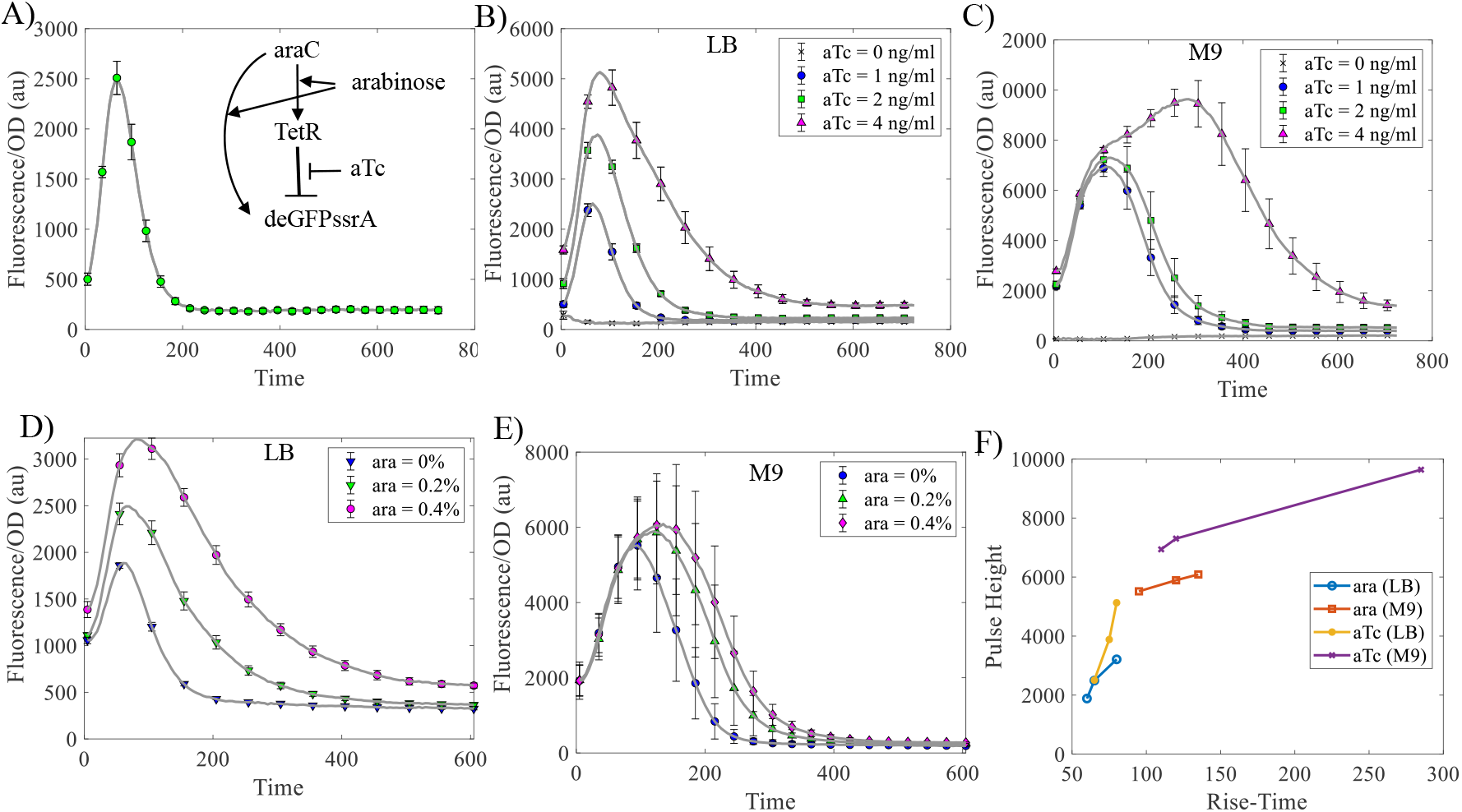
Experimentally obtained pulse trajectories for different levels of inducers. A) Solid line represents the pulse generated by incoherent feedforward loop (inset) for *aTc* = 1 *ng/mL* and *arabinose* = 0.2 %. Error bar represents standard deviation of three separate repeats. Pulse trajectories for *aTc* variations at 0.2 % *arabinose* in B) LB media and C) M9 media. Pulse trajectories for *arabinose* variations at 2 *ng/mL aTc* for D)LB media and E) minimal media. F) Solid lines with indicated symbols represent the pulse height and rise time co-variations for above description.

We used the inducers to experimentally explore the parameter space in terms of the properties of the pulse. We repeated the experiment for different *arabinose* levels at a fixed *aTc* level, and for different *aTc* levels at a fixed *arabinose* level (Fig. 3B-E). We find that as *arabinose* levels increase pulse height increases and rise time increases. Further, we find that as *aTc* levels increase, the pulse height increases and rise time increases. These results shows that for these experimental conditions, there is a constraint that as the pulse height increases, the rise time increases, making a higher amplitude pulse also slower (Fig. 3F).

### Systematic Model Analysis

While we find, experimentally, that as the pulse height increases, the rise time also increases, these results do not match the expectations from the standard model considered above. In the model, an increase in *arabinose* should change the parameter dissociation constant of *AraC* from *K*_*u*0_ to *K*_*u*_ (*K*_*u*_ > *K*_*u*0_). As this parameter increases, we find the pulse height decreases and rise time increases (Fig. 4A). This is not seen experimentally. Similarly, an increase in *aTc* should change the parameter dissociation constant of *TetR* from *K*_*x*0_ to *K_x_* (*K*_*x*0_ > *K_x_*). As this parameter increases, we find the pulse height increases with no change in the rise time (Fig. 4A). This is also not seen experimentally. A possible reason for this discrepancy could be the difference in experimental circuit in relation to the model.

**Figure 4:**
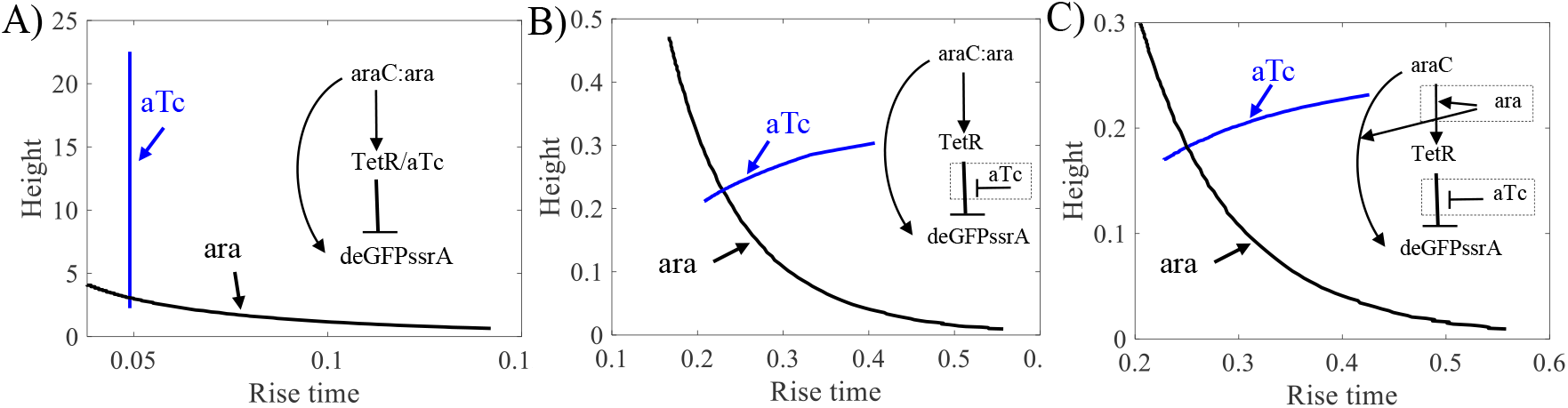
Amplitude-rise time co-variation for change of the inducers in computational models A) Solid lines represent the amplitude and rise time co-variations in the standard model for inducers. Indicated with arrow for aTc and ara (*arabinose*). B) Solid lines represent the amplitude and rise time co-variations in the modified model of *aTc − TetR* dynamics for inducers. Inset figure represents the modification in the model compared to inset figure in A. C) Solid lines represent the amplitude and rise time co-variations in the modified model of *aTc − TetR* and *arabinose − AraC* dynamics for inducers. Inset figure represent the modification in the model compared to inset figure of A.

To investigate these discrepancies, we first expand the model to include *TetR-aTc* interactions. An assumption used in the standard model is that the repression is strong. Therefore, we replaced the term 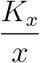 with a general term for repression 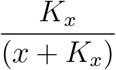 to obtain a modified model (see Methods). We find that the modified model can replicate the experimental results as far as *aTc* is concerned (Fig. 4B). To understand how the modified model captures the experimental results, we considered the analytical solution of the models. For the standard model, with equal degradation rate parameters, the output solution is,

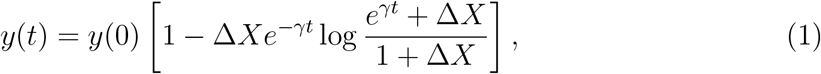

where 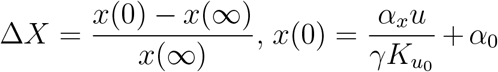, and 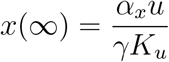. This is independent of the dissociation constant *K_x_* for the standard model. The solution of the modified model is,

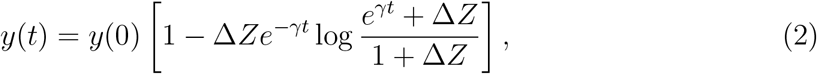

where 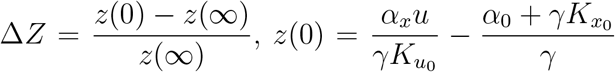, and 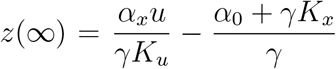. We note that the solution of modified model depends on *K_x_* which is not true for standard model. Hence, the timescales in the standard model are unaffected by perturbation in *K_x_*.

Next, we similarly expanded the model to include *arabinose-AraC* dynamics (see methods). We replaced the term 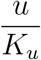 with a general term for activation 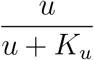 However, we could not replicate experimental results (Fig. 4C). The output solution of this model is,

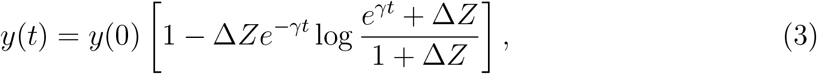

where 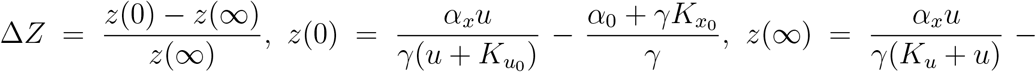 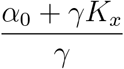. The solution has similar timescale dependency as the earlier models. Therefore, the model is not able to capture the experimental results when *arabinose* is changed, just like in the earlier models. This is a gap in our understanding of the experimental results in relating to the model. The gap could be due to aspects such as ignoring the resources needed to produce proteins and other possible dynamics such as host-circuit interactions or resource competition.

## Summary

Understanding constraints in the co-variation of amplitude and timescale in a biomolecular pulsing circuits is important for their design. Using experimental measurements and mathematical models of a benchmark pulse generating biomolecular circuit — the incoherent feedfoward loop — we address this issue and present three main points. First, we explore the parameter space of a widely used model of an incoherent feedforward loop and find global trends for the covariation of amplitude and rise time. Second, we find experimental evidence, that as amplitude of the pulse increases, the rise time also inceases. Third, we discuss the inconsistencies between the mathematical model and experimental measurements and ways to resolve these. These results provide experimental evidence for the existence of constraints in the design space of such pulse generating circuits.

## Methods

### Materials

The plasmid for feedforward loop is pBEST-OR2-OR1-Pr-araC, pBAD-TetR, pBADTetO1-deGFP-ssrA was from plasmids from the lab of Prof. Richard M. Murray (Addgene plasmid # 45789; http://n2t.net/addgene:45789; RRID:Addgene_45789). This was transformed into the *E. coli* MG1655 strain background. The realization is based on transcriptional activator *AraC*, transcriptional repressor *TetR*, and reporter *deGFPssrA* as in Fig. 3A. The *deGFPssrA* is a green fluorescent protein with ssrA degradation tag at C-terminal of *deGFP* protein. The protein *AraC* is constitutively expressed with promoter *Pr-OR2-OR1*. The *TetR* and *deGFP* are expressed with promoter *P_BAD_* that can be activated by *AraC* in presence of arabinose. Additionally, the operator sites for *TetR* are fused to a promoter *P_BAD_* regulating the *deGFP*. This allows *TetR* to repress the promoter. The circuit plasmid has Ampicillin resistance marker and ColE1 origin of replication. All these three genes are on the same plasmid. The untranslated region for all of this is UTR1, which has a strong ribosome binding site. The transcriptional terminator for all three is called T500. In 5’ to 3’ order, the genes are: pBAD-tetO1 (repressed by TetR)-UTR1-deGFP-ssrA-T500, pBAD-UTR1-TetR-T500, and OR2-OR1-Pr (bacteriophage lambda with one mutation)-UTR1-araC-T500. The pBAD-TetR insert is inverted relative to reporter gene.

### Measurement Protocol

For experimental measurement, the strain was inoculated in LB media supplemented with Ampicillin for 16 hours. The culture was subsequently diluted 1:200 in fresh minimal media and LB media supplemented with Ampicillin. The culture was induced with *aTc* (1, 2, 4 *ng/ml*) and incubated for two hours. Next, different levels of *arabinose* (0, 0.01, 0.02, 0.04%) was added to the incubated culture and this final culture was used for plate reader measurement. The measurement was taken for ten hours at 37 °*C* with five minute sampling and two minute shaking between successive readings. The samples were placed in triplicate and the above protocol was repeated for three days.

### Data Analysis

MATLAB was used for data analysis. A blank sample containing only media and no cells was used as background and subtracted from culture reading for both optical density and fluorescence.

### Model Simulations

The standard mathematical model of the incoherent feedforward model is [13],

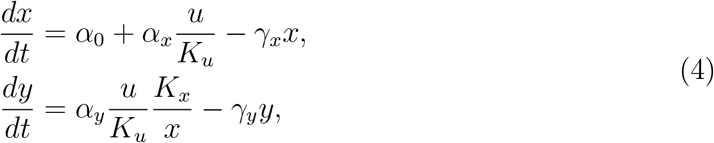

where *u* represents *AraC:arabinose* complex act as input, *x* represents the ‘free’ *TetR*, the output *y* represents *deGFPssrA*. The model parameters *γ_x_* is the degradation rate for protein *x*, *γ_y_* is the degradation rate for protein *y*, *α_x_* is the production rate of protein *x*, *α_y_* is the production rate of protein *y*, and *K_x_* is the dissociation constant for the binding of *x* to the promoter of *y*. For simulation, the parameters values considered are *α*_0_ = 0.01 *nM/hr*, *α_x_* = 10 *nM/hr*, *α_y_* = 10 *nM/hr*, *γ_x_* = 1 1/*hr*, *γ_y_* = 10 1/*hr*. The model assumes only strong activation from *u* and strong repression from *x*, hence the approximated term and 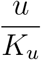 and 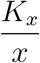. More generally, repression can be represented by 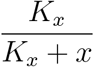 instead of 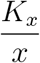, so that modified model becomes,

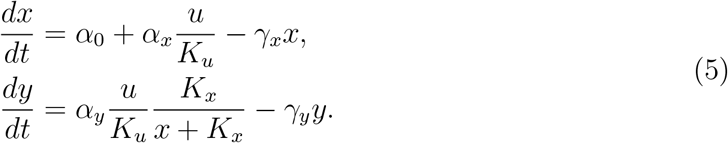

Similarly, a more general activation is modelled as 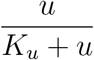 instead of 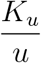,

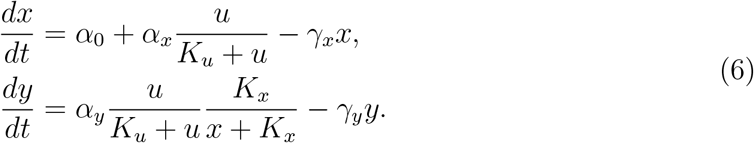

These models are simulated in MATLAB using the function *ode45* with default settings.

For Fig. 2A, individual parameters (*α_x_*, *α_y_*, *γ_x_*, *γ_y_*) are varied from 0.1 to 10. For random simulation, individually varied Fig. 2B and simulaneously varied Fig 2C, parameter sets are uniformly sampled from *α_x_* = 1 − 100, *α_y_* = 1 − 100, *γ_x_* = 0.1 − 10, *γ_y_* = 0.1 − 10 using *random* function from MATLAB.

## Acknowledgement

The authors would like to thank Richard Murray for the gift of pBEST-OR2-OR1-Pr-araC, pBAD-TetR, pBAD-TetO1-deGFP-ssrA (Addgene plasmid 45789).

